# Heartbeat evokes a dynamically changing cortical theta-synchronized network in the resting state

**DOI:** 10.1101/524108

**Authors:** Jaejoong Kim, Bumseok Jeong

## Abstract

In the resting state, heartbeats evoke cortical responses called heartbeat-evoked responses (HERs). While previous studies reported regional level HERs, researchers have not determined how heartbeat is processed at the cortical network level. Using resting-state magnetoencephalography data from 87 human subjects of both genders provided by the Human Connectome Project, we first showed that heartbeat increases the phase synchronization between cortical regions in the theta frequency, which forms a network structure, and we called this network a heartbeat-evoked network (HEN). The HEN was not an artefactual increase in phase synchronization. The HEN was partitioned into three modules with connector hubs in each module. The first module contained major interoception-related regions and thus was called a visceromotor-interoceptive network (VIN) displaying the strongest synchronization among modules, suggesting a major role for the VIN in processing heartbeat information. Two modules contained regions involved in the default mode network (DMN). The HEN structure was not fixed, but dynamically changed. The most prominent change was observed at approximately 200 ms after R-peak of the electrocardiogram, which was quantified based on the ‘flexibility’ of the network. Furthermore, the strongest synchronization within VIN was observed before heartbeat stimulated the cortex, which might be related to the prediction of an afferent heartbeat signal, thus supporting an interoceptive coding framework. Based on our results, the heartbeat is processed at the network level, and this result provides a useful framework that may potentially explain previous results of the regional level HER modulation through network-level processing.

**Significance statement:** The resting-state network is composed of several networks supporting different functions. However, although the heartbeat is processed in the cortical regions, even in the resting state, the network supporting this function is unknown. Thus, we identified and investigated the heartbeat-evoked network (HEN), a network composed of significantly increased theta-phase synchronization between cortical regions after a heartbeat. The HEN comprised three modules. In particularly, the visceromotor-interoceptive network was likely to play a major role in network-level heartbeat processing and displayed the strongest synchronization immediately before the heartbeat enters the CNS, which supports an interoceptive predictive coding framework. These results provide a novel framework that may improve our understanding of cortical heartbeat processing from a network perspective.

## Introduction

In the resting state, brain networks supporting diverse functions exist and are called resting-state networks (RSNs) (Thomas Yeo et al., 2011); these networks interact with each other (Ashley C Chen PNAS 2009). RSNs were recently shown to be closely related to the intrinsic functional network architecture of the brain that shapes the brain network structure during various tasks (Cole, Bassett, Power, Braver, & Petersen, 2014). Therefore, an understanding of the RSNs, including how they are classified into networks with different functions and which region plays an important role is important for understanding the intrinsic network architecture that occurs across most brain states (Cole et al., 2014).

One of the fundamental, but less frequently investigated, RSNs is the network that processes the signals from the viscera. In particular, because the heartbeat signal and the signal induced by gastrointestinal pacemaker cells continuously enter the central nervous system, even in the resting state (Tsakiris & De Preester, 2018), RSNs processing these visceral signals likely exist. Indeed, a recent study revealed the existence of the Gastric Network, whose BOLD time series are phase synchronized with the basal gastric rhythm (Rebollo, Devauchelle, Béranger, & Tallon-Baudry, 2018). However, the RSN processing the heartbeat signal has not been investigated. In this study, we have investigated the RSN evoked by the heartbeat.

The heartbeat signal continuously enters the brain and evokes cortical activity, termed heartbeat-evoked responses (HERs) (Pollatos & Schandry, 2004). This entry of the heartbeat into brain is mainly mediated by baroreceptor stimulation, which occurs approximately 200 ms after cardiac contraction (Eckberg & Sleight, 1992), and the HER effect is known to occur between 200-650 ms after R-peak (Kern, Aertsen, Schulze-Bonhage, & Ball, 2013). Previous studies reported HERs in various regions, including the right insula and the anterior cingulate cortex during the heartbeat perception task and emotion processing (Couto et al., 2015; Pollatos, Kirsch, & Schandry, 2005), the posterior cingulate cortex and supplementary motor area in the bodily self-consciousness modulation task (Park et al., 2016), and the precuneus and ventromedial prefrontal cortex in the mind wandering task (Babo-Rebelo, Richter, & Tallon-Baudry, 2016; Babo-Rebelo, Wolpert, Adam, Hasboun, & Tallon-Baudry, 2016).

While previous studies have reported a region-level HER effect, the network-level explanation of the HER effect, namely, how brain regions are functionally connected by the heartbeat to form a network, remains unknown. We hypothesized that heartbeat would evoke a functional coupling between brain regions that form a network structure in the resting state. Using the resting-state MEG dataset, we have investigated the heartbeat-evoked network (HEN), which was defined as a network composed of significantly increased phase synchronization between regions compared with baseline values, to test this hypothesis. We first showed the existence of the HEN and then investigated the structure of the HEN. In particular, because HERs evoked during different tasks involve different regions, we hypothesize that the HEN, which is an intrinsic network evoked by heartbeat, is partitioned into several subnetworks called ‘modules’ that might supports different functions and would have hubs connecting each module, enabling effective interactions between modules (Fornito, Zalesky, & Bullmore, 2016). This hypothesis was tested using the modularity analysis (Bassett et al., 2011).

Furthermore, inspired by the interoceptive predictive coding framework, which assumes the top-down prediction and the bottom-up processing of interoceptive signals (Barrett & Simmons, 2015), we expected that the HEN structure would differ when predicting heartbeat and performing bottom-up processing of heartbeat signals, and the changes in the two structures would occur at the time the heartbeat stimulates the cortex, which is approximately 200ms after R-peak. We performed a dynamic modularity analysis to test this hypothesis and showed the changing network structure over time; we also investigated when and how this network change occurs (Bola & Sabel, 2015). Finally, we hypothesized that the HEN would support the interoception-related function, which was tested by determining relationship between the HEN and the network regulating emotion perception.

## Methods

### Dataset description

Resting-state MEG data from 89 subjects collected from the human connectome project (HCP) S1200 data release were used in this study (Larson-Prior et al., 2013; Van Essen et al., 2013), and all subjects were young (22-35 years of age) and healthy. MEG recordings were collected on a whole-head Magnes 3600 scanner (4D Neuroimaging, San Diego, CA, USA) with 248 magnetometer channels at a sampling rate of 2034.51 Hz. Recordings were performed in three sessions, and each session lasted 6 minutes. HERs were extracted from the preprocessed version of the MEG dataset, which is publicly available at Connectome DB (Hodge et al., 2016). The preprocessing pipeline of HCP data included segmentation of the raw data into epochs of 2 seconds and the removing bad segments and bad channels. Importantly, an independent component analysis (ICA) (Hämäläinen, Hari, Ilmoniemi, Knuutila, & Lounasmaa, 1993) was applied to remove the cardiac field artifacts (CFA) and electrooculography (EOG)-related artifacts, and data were finally downsampled to 508.68 Hz. Notably, because these preprocessed data did not include electrocardiogram (ECG) recordings, which are essential for HER extraction, we used an ECG recording included in the raw MEG data. Among 89 subjects, two subjects were excluded (IDs 149741 and 17746) because we failed to detect the R-peak in the ECG recording of subject 149741. In subject 177741, an error occurred when performing automated anatomical labeling (AAL) atlas-based source time course extraction (the time course extraction of ‘Occipital_Sup_L’ failed in this subject). Finally, 87 subjects were included in our analysis (47 males and 40 females).

## MEG analysis

### HER extraction procedure

HER extraction and preprocessing were performed using the FieldTrip toolbox (Oostenveld, Fries, Maris, & Schoffelen, 2011). First, preprocessed HCP MEG data, which were initially segmented into 2 second epochs, were concatenated into one continuous time series such that each segment was realigned to its original location in the raw MEG recording (preprocessed data contained information about the locations of each segment in the raw MEG recording). Because the bad segments were removed, the concatenated continuous time series data had empty spaces where bad segments existed. These empty spaces were replaced by NaN. Next, the R-peak was detected in the ECG recordings using the Pan-Tomkins algorithm (Pan & Tompkins, 1985). Then, epoching of HERs from 900 ms before R-peak to 1800 ms after R-peak was performed on the concatenated continuous MEG data. As mentioned above, the concatenated MEG data contained the time window including NaN; thus, the HER epoching procedure resulted in some NaN-containing epochs. These NaN-containing HER epochs were removed. Finally, the HER extraction procedure resulted in 838.5 (± 137.8) HER epochs on average. The mean interbeat interval (IBI) of every subject was 983.0 ms (± 150.2, corresponding heartrate: 61.03 beats per minute (BPM)) with a range of 675.4-1375.2 ms (heartrate range: 43.63-88.84 BPM).

### Source reconstruction of HERs

Every HER data sensor was source-reconstructed using the linearly constrained minimum variance (LCMV) beamformer methods (Van Veen, Van Drongelen, Yuchtman, & Suzuki, 1997) provided in the FieldTrip toolbox in a manner similar to that used in a previous study (Heusser, Poeppel, Ezzyat, & Davachi, 2016). A common spatial filter was estimated for each source point using HER data from all trials, an HCP-provided single-shell volume conduction head model and an HCP-provided 4 mm grid source model for every subject (Larson-Prior et al., 2013). Then, this common spatial filter was applied to sensor HER data (sensor * time matrices) to calculate the time courses of each source. Finally, we used AAL atlas-based parcellation (Tzourio-Mazoyer et al., 2002) to perform a region of interest (ROI)-based connectivity analysis. Among the AAL atlas regions, we only used 78 cortical regions after excluding subcortical and cerebellar regions, and the time courses of each region were averaged (Supplementary Table 1). This final step produced the time courses of HERs for every region, epoch and subject.

### Calculation of the phase locking value in the theta frequency range between cortical regions

The phase locking value (PLV) (Lachaux, Rodriguez, Martinerie, & Varela, 1999) was used as a measure of functional connectivity between the 78 cortical regions. The PLV between two regions, x and y, was defined as follows:

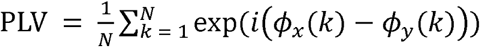

where N is the number of trials, k is the index of each trial and *ϕ* is the phase. The consistency of the phase lag between x and y and ranges from 0 to 1. The PLV was calculated in the steps described below using the functions of the FieldTrip toolbox. We hypothesized that synchronizations would occur in the theta band (4-7 Hz), the frequency band with the strongest increase in phase synchronization within regions induced by heartbeat in a previous study (Park et al., 2017). First, complex Morlet wavelet transformation was performed on every trial with a 20 ms time step from −300 ms to 600 ms R-peak and a frequency ranging from 4 to 7 Hz with 1 Hz steps. The number of cycles used in the wavelet transformation was 4. Then, the PLV was calculated for every pair of regions in each time and frequency step. PLVs from 4 to 7 Hz were averaged to obtain the PLV of the theta frequency range. These procedures resulted 78 (number of ROIs) by 78 by 31 (time windows from −300 to 600 ms R-peak with 20 ms steps) PLV matrices for each subject.

### Identification of the static HEN using network-based statistics

We compared the PLVs between the baseline period, which was defined as a time window before 300 ms to 100 ms R-peak onset, and the time window after 200 ms to 600 ms R-peak onset, which is the time window in which the effects of HERs were reported in most previous HER studies (Fukushima, Terasawa, & Umeda, 2011; Pollatos & Schandry, 2004), to determine whether the heartbeat evoked a network composed of significantly increased phase synchronization between regions, and we called this time window the ‘evoked’ period. Notably, the 200 ms period after R-peak corresponds to the approximate time that the heartbeat signal enters the CNS following carotid baroreceptor stimulation (Eckberg & Sleight, 1992) and the baseline period used in the present study is the period used in a previous study of HER-evoked phase synchronization within regions. This baseline period was postulated to avoid cardiac artefacts around the ECG P-wave (Park et al., 2017).

We then performed a group-level network-based statistical (NBS) (Zalesky, Fornito, & Bullmore, 2010) analysis, a statistical method that controls multiple comparisons at the network level. This analysis enabled us to identify a network composed of significantly increased PLVs between cortical regions in the evoked period compared to the baseline period at the group level. First, baseline and evoked PLV matrices were computed by averaging PLVs from each time window for every subject, which resulted in one baseline PLV matrix and one evoked PLV matrix for each subject. Second, multiple paired t-tests comparing PLVs from the evoked period and the baseline PLV were performed for every pair of cortical regions, which resulted one matrix of t-values from these paired t-tests. Then, a threshold t-value of 1.99, which corresponds to a p-value of 0.05 at 86 degrees of freedom, was applied to the matrix of t-values, and thus a t-value less than 1.99 was set to 0. The network statistic was computed by adding the t-values of all the connected components in the thresholded matrix (connected component means that any two nodes within this components are connected by a path of edges). Next, a null distribution of the network statistic was created from 5000 permutations by randomly permuting an element of the evoked PLV matrices and the baseline PLV matrices within each subject. Finally, network-level familywise-error (FWE)-corrected p-values of the network were obtained using the original network statistic and null distribution. Next, we constructed a heartbeat-evoked synchronization (HES) matrix whose elements corresponded to the increase in PLV in the evoked period compared to the baseline period, and each element was significant in the NBS results. Therefore, the HES matrix was composed of elements with significantly increased PLVs in the group-level NBS and represents the structure of the HEN. Notably, the current network analysis did not consider the dynamic changes in the HEN across the 200-600 ms period; thus, we called the HEN identified in the present study a ‘static’ HEN to discriminate it from the ‘dynamic’ HEN, which will be described later.

### Examination of increased theta phase synchronization between ECG signals and brain regions

We postulated that the static HEN we identified might represent an artificial increase in phase synchronization caused by a CFA. We expected that if an electromagnetic field induced by cardiac contractile activity directly influenced both region A and B and this effect artificially increased phase synchronization between these two regions, the phase synchronization would have increased between regions A and B and the phase synchronization between the ECG signal and both regions A and B should have increased after a heartbeat, because the same electromagnetic field induced by cardiac contraction influenced all three signals, including the ECG signal and the signals from regions A and B. We assessed whether theta phase synchronizations between ECG signals and brain regions increased during an evoked period (200-600 ms post R-peak) compared to the baseline to test this hypothesis. We calculated the PLVs between ECG signals and 78 cortical regions in the theta band for every subject, which resulted two 78 by 1 vector of ECG-brain region PLVs from the baseline and evoked period for every subject. Then, we performed 78 group-level paired t-tests between the PLVs from the evoked and baseline periods for all 78 cortical regions to determine which PLVs between each ECG-brain region pair were significantly increased in the evoked period compared to baseline.

### Surrogate R-peak analysis

As another control analysis, we tested whether the HEN was time-locked to the heartbeat. We created 100 surrogate R-peaks that were independent of the original heartbeats (Babo-Rebelo, Richter, et al., 2016; Park et al., 2016; Park, Correia, Ducorps, & Tallon-Baudry, 2014). Surrogate R-peaks were created by randomly shifting the original R-peaks (−500 ms ∼ +500 ms) by the same amount in each subject. Then, we computed the surrogate HERs with surrogate R-peaks, and the PLV of the theta band was subsequently calculated for each set of surrogate R-peaks. Finally, an NBS analysis comparing the average PLV after the 200-600 ms surrogate R-peak with the baseline PLV (300 ms to 100 ms before the surrogate R-peak) was performed for each set of surrogate R-peaks. Finally, the surrogate distribution of the maximum network statistics of the surrogate R-peaks was determined, and the p-value of the original network statistic within the surrogate distribution was calculated.

### Identification of the modular structure within the static HEN

We applied a community detection algorithm to the HES matrix to determine how the static HEN is partitioned into different subnetworks. Optimal partitioning of cortical regions was performed using the Louvain greedy algorithm (Blondel, Guillaume, Lambiotte, & Lefebvre, 2008) to maximize the modularity index Q formulated using the following equation:

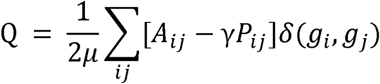

In this equation, *A*_*ij*_ represents the strength of the edge between node i and node j, *P*_*ij*_ represents the expected weight between node i and node j, *µ* is the sum of the strengths of all edges in the network, and *δ*(*g*_*i*_ *g*_*j*_) is 1 if node i and j belong to the same community and 0 otherwise (*g*_*i*_ is a label of the community to which node i belongs). The resolution parameter *γ* was set to 1, which is a default value. However, because the partition that maximizes Q can vary across each algorithm run, we used the consensus partition method to identify the most representative partition S (Lancichinetti & Fortunato, 2012) using the functions of the Brain Connectivity toolbox (BCT) (Rubinov & Sporns, 2010). The consensus partition procedure, which is identical to a previously reported procedure (Fornito et al., 2016), is briefly explained below. First, a community detection algorithm (Louvain greedy algorithm) was run 10000 times to create 10000 partitions. Second, the agreement matrix D was constructed. Each element of D corresponded to the proportion of the number of times that nodes i and j were in the same module to the number of total iterations. Third, a threshold τ = 0.2 was applied to D. The value of τ was chosen to be less than 0.4, which was recommended in a previous study (Lancichinetti & Fortunato, 2012). Fourth, community detection was performed 10000 times using D, which created another agreement matrix, D’.

Fifth, steps 2 through 4 were repeated until the consensus matrix exhibited a block-diagonal structure in which all edge weights equaled one for node pairs in the same community and zero otherwise. We initially constructed the agreement matrix with 10000 iterations of the HES matrix. Then, 10000 partitions were provided as the functional input for steps 2 through 4, and these processes were repeated until convergence was achieved. One-way analysis of variance (ANOVA) between the average edge t-values of three modules was performed to compare the mean synchronization strengths of each module. Subsequent post hoc two-sample t-tests were also performed.

### Identification of the hubs of the static HEN

One of the important features of a network is the hub of the network, which is defined as a node that plays an important role within the network, such as connecting nodes within network (Fornito et al., 2016). Specifically, we hypothesized that hubs that connect modules of the static HEN exist and enable effective interactions between modules. This type of hub is called a ‘connector’ hub and is defined by the graphical theoretical measurements called within-module degree z-score (Guimera 2005) and the participation coefficient (Guimera 2005). The within-module degree z-score was calculated using the following equation:

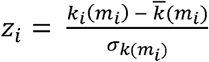

where *k*_*i*_(*m*_*i*_)is the within-module degree (strength), 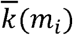 is the mean within-module degree (strength) of all nodes in the same module as node i, and σ_*k*_(*m*_*i*_) is the standard deviation of *k*_*i*_(*m*_*i*_) values across all nodes in module *m*_*i*_. This measure quantifies a normalized within-module strength and was calculated using the BCT function ‘module_degree_zscore.m’. Finally, the participation coefficient quantifies the nodes participation to each module as follows:

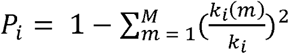

where M is the number of modules in the network and *k*_*i*_(*m*) is the strength of the edge between node i and other nodes in module m. Using within-module degree z-scores and participation coefficients, we defined role of every node according to the z-P classification (Guimera & Amaral, 2005). In particular, the connector hub is a node with many connections within the module to which the node belongs and also forms many connections with nodes of other modules; thus, the connector hub efficiently connects nodes within one module to other modules (Fornito et al., 2016). In our study, the connector hub was defined as a node with a within-module degree z-score greater than 2.5 and a participation coefficient (P) greater than 0.3 (Fornito et al., 2016; Guimera & Amaral, 2005).

Additionally, other types of hubs in the static HEN displayed high betweenness centrality, which is defined as the largest fraction of all the shortest paths in the network that passes through a given hub (Brandes, 2001), and the hubs with high strength - namely, the hubs whose sums of the weights of all edges connected to that hub were high - were calculated. The graph theoretical measures defining hubs were calculated using the functions of the BCT.

### Analysis of the dynamic HEN

We a performed dynamic modularity analysis to test how the HEN structure changes over time (Bassett et al., 2011). In contrast to the modularity analysis performed for the static HEN defined by increased average phase synchronization in 200-600 ms post R-peak compared to the baseline (300-100 ms before R-peak), we investigated how the HEN defined in a shorter time scale of 20 ms evolves over time, and the assembled short time scale HENs over the whole time window was called the ‘dynamic’ HEN. We hypothesized that structural changes occurring within the dynamic HEN over time were related to 1) the composition of each HEN module (which regions comprise each network), 2) the interaction pattern within nodes contained in each network and 3) the interaction between networks. We hypothesized that the maximum changes within the dynamic HEN structure would occur at approximately 200 ms post R-peak because the heartbeat is thought to stimulate the cortex at approximately that time point (Eckberg & Sleight, 1992). We extended the time window of our analysis from 200-600 ms to 0-600 ms post R-peak, the time window that starts from the onset of R-peak, to compare the changes before and after 200 ms post R-peak.

### Detection of the changing subnetwork structure within the dynamic HEN

The changing module structures of the dynamic HEN were computed using a multilayer modularity analysis (Mucha, Richardson, Macon, Porter, & Onnela, 2010). In this analysis, each layer contains the graph structure of each time window (which, in our analysis, was the HES matrix of each time point). Similar to the detection of the static community structure, multilayer partitioning maximizes the multilayer modularity index *Q*_*ML*_ by considering temporally linked community structures, meaning that the algorithm assumes that the region belongs to community A at time t is likely to belongs to the same community at time t+1, which is formulated as

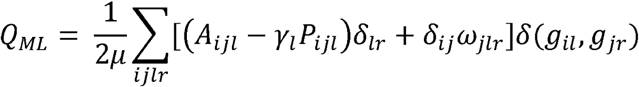

where the indexes *l* and *r* represent a layer (the layer represents the time window). *A*_*ijl*_,*P*_*ijl*_, *γ*_*l*_, and *µ* are defined in a similar manner to the definitions used in the static modularity analysis. The term (*A*_*ijl*_- *γ*_*l*_ *P*_*ijl*_)*δ*_*lr*_ considers the coupling within the layer where *δ*_*lr*_ = 1 if *l* = *r*. Therefore, similar to the static modularity calculation, (*A*_*ijl*_ - *γ*_*l*_ *P*_*ijl*_) is calculated within single layers. The term *δ*_*ij*_ω_*j;r*_ reflects the coupling between layers, where ω_*jlr*_ is an interlayer coupling parameter between layers *l* and *r* for node j and is typically the same for every *jlr* (called ω hereafter). If ωis large, the number of changes in the community assignment across layers decreases, while small *ω* (< 1) values increase the number of changes. While a single established optimal parameter does not exist, the typical value selected for *ω* is 1 (Bassett et al., 2011; Telesford et al., 2016). However, we hypothesized that only a slight change would occur during the time course because the HEN, which is an intrinsic brain network, should have a stable structure with little flexibility relative to other exteroceptive task-driven networks. Therefore, we set *<* to 0.1, which is the lowest value that was tested in a previous study (Bassett et al., 2013), allowing us to investigate the changing network structure over time. Moreover, we tested multiple values of < from 0.1 to 1.05 with a step size of 0.05 and determined that no community assignment changes across layers occurred at < values greater than 0.2. *δ*_*ij*_ equals 1 if node i = j and equals 0 otherwise. Finally, the term δ (*g*_*il*_, *g*_*jr*_) equals 1 only when node i in layer *l* and node j in layer *r* are in the same community and equals 0 otherwise. Therefore, *Q*_*ML*_ is maximized both by maximizing the intralayer modular structure and by maximizing the interlayer coupling of the same node.

We constructed a multilayer network using the steps described below. First, the 31 HES matrices of the time windows starting from 0-20 ms post R-peak to 580-600 ms post R-peak were constructed, where the HES matrix of each time window was constructed in the same way as described in the static HEN analysis by performing an NBS analysis between the PLV matrices of each time step and the baseline period (300-100 ms before R-peak, which is same as the static HEN analysis). Then, the resulting 31 HES matrices were combined to construct a multilayer network, which represents the dynamic HEN. Because we only used elements (increased PLV compared with the baseline) with p-values less than 0.0002 (network-level FWE-corrected) at all 31 time points, the surviving networks of each layer also had p-values less than 0.01, even when the Bonferroni correction was applied to the p-values of all 31 time points (the corresponding p-value was less than 0.0062). The dynamic HEN was partitioned using the Louvain-like locally greedy algorithm for multilayer modularity optimization (Mucha et al., 2010). Similar to the method used in the static modularity analysis, we applied the consensus partition method to obtain the representative multilayer partition *S*_*ml*_ (Braun et al., 2015). We computed multilayer partitions 10000 times to ensure that the matrices of each layer agreed. Then, the same consensus partition method that was used in the static modularity analysis was applied to each layer, resulting in consensus partitions for every layer. Finally, community labels of every layer were re-labeled in a way that maximized the persistence of communities using the ‘multislice_pair_labeling.m’ function (Bazzi et al.).

### Flexible changes in module composition over time

We hypothesized that the composition of the HEN modules would change over time, and this change would be maximized at approximately 200 ms post R-peak. We calculated an index called ‘flexibility’ to quantify the degree of change (Bassett et al., 2011). Flexibilities of the entire brain at each time step (F_time) were calculated using the optimal partition *S*_*ml*_, which was an output of the consensus partition of the multilayer network (Braun et al., 2015). The flexibility of the entire brain (F_time) at one layer (time point) represents the proportion of regions that changed their community assignment between two consecutive layers (between layer k and k+1) (Braun et al., 2015). The F_time was calculated for each time step from 0 ms to 580 ms post R-peak, thus resulting in 30 values for F_time.

### Dynamic changes in synchronization between heartbeat-evoked networks

Consensus partitioning of the dynamic HEN resulted in three HEN modules that changed over time. These modules were matched to three modules identified in the static HEN. Next, we tested how the within- and between-module synchronizations of each module changed over time. We first divided time window from 0-600 ms post R-peak into equally sized 3 time bins, namely, 0-200 ms, 200-400 ms, and 400-600 ms post R-peak to statistically compare changes in the within- and between-module synchronizations over time. The sizes of these bins were determined to 1) enable comparisons between 0-200 ms and later time windows and 2) ensure equally sized time bins equal. Then, we computed the within- and between-module synchronizations for every subject by creating a phase locking value difference matrix (PDM) with a size of 78 (number of cortical regions) by 78 by 31 (number of time steps from 0-600 ms post R-peak) for every subject. The elements of the PDM at time step k represented the difference between the PLV at time k and the baseline value, and the elements that did not belong to the dynamic HEN were set to 0. Then, the within- and between-module connectivities were calculated using the PDMs. All intracommunity elements of PDM were averaged at each time point to calculate the within-module synchronization in each subject. The between-module synchronizations at each time point were calculated by averaging the elements of PDM reflecting the PLV between the regions of two different modules. Because three HEN modules were identified, these procedure resulted in three within-module synchronization values and three between-module synchronization values for the three time bins for every subject. Then, we statistically compared the within- and between-module synchronizations in these time ranges. Six (three within-module synchronizations and three between-module synchronizations) separate 1 by 3 (time bins) one-way repeated measures ANOVAs were performed for the within/between-module synchronizations of the three time windows. Post hoc t-tests were also performed for each test.

### Relationship between the emotion recognition abilities of the subjects and the within/between-module synchronizations of the HEN

The features of RSNs are related to the performance of tasks assessing working memory and executive function (Reineberg, Andrews-Hanna, Depue, Friedman, & Banich, 2015; van Dam, Decker, Durbin, Vendemia, & Desai, 2015). Similarly, we hypothesized that the features of the HEN would support the cardiac interoception-related functions, such as emotion perception, which was shown to modulate the HERs in previous studies employing the emotion perception task (Couto et al., 2015). We performed a correlation analysis between network features of the HEN and the abilities of subjects to recognize the emotions associated with fearful and angry faces, which are particularly known for their relationships with cardiac interoception (Garfinkel & Critchley, 2016; Marshall, Gentsch, Jelinčić, & Schütz-Bosbach, 2017), to test this hypothesis. We used score of the Penn Emotion Recognition Test for anger and fear (ER40ANG, ER40FEAR) to quantify the abilities of the subjects to recognize the emotions associated with fearful and angry faces (Gur et al., 2001), which was included in the HCP S1200 dataset. Briefly, in this test, participants were sequentially presented 40 faces and asked to choose the emotion the face was showing from among five choices: happy, sad, angry, scared and no feeling. Furthermore, the network features we used here were the time courses of the within- and between-module synchronizations for every subject, where within-module synchronization represents the feature of each module and the between-module synchronization represents the feature related to interactions between modules. The between-subject correlation analyses between the emotion recognition score and module synchronizations were performed at each time point, and multiple comparisons for every time point multiplied by six synchronizations (three within-module synchronizations and three between-module synchronizations) were controlled using the cluster-based permutation correlation test (Oostenveld et al., 2011). Briefly, in the cluster-based permutation correlation test, similar to NBS, the cluster correlation statistic was calculated by summing all the t-values of the cluster at significant time points (thresholded using an uncorrected p-value < 0.05 for the initial correlation analysis). Then, 5000 permutations were performed, and the maximum cluster t statistic was extracted for each permutation to generate a null distribution. Finally, the original cluster statistic was evaluated for this null distribution.

## Results

### Phase synchronizations between cortical regions increased at 200-600 ms post R-peak, confirming the existence of the HEN

Before performing the statistical test using NBS, we found that among the total of 78C2 pairs of cortical regions, 60.8% showed a positive PLV (Figure 1a) in the evoked period compared with the baseline, while other 39.2% showed a negative PLV compared with the baseline (Figure 1b). These results indicates that heartbeat induces both an increase and decrease in the phase synchronization between regions; however, our interest was the network displaying a statistically significant increase in phase synchronization. The NBS analysis statistically confirmed a network displaying a significant increase in phase synchronizations in the evoked period compared to the baseline (network-level FWE-corrected p < 0.001), which revealed the existence of the static HEN. The density of the network was 14.89%, and the phase synchronization between ‘Cingulum_Mid_R’ and ‘Olfactory_L’ had the highest t-value among the regions (t (86) = 6.67, Figure 1a). Representative plots of the decreased phase synchronizations between regions are shown in Figure 1b.

**Figure 1.**
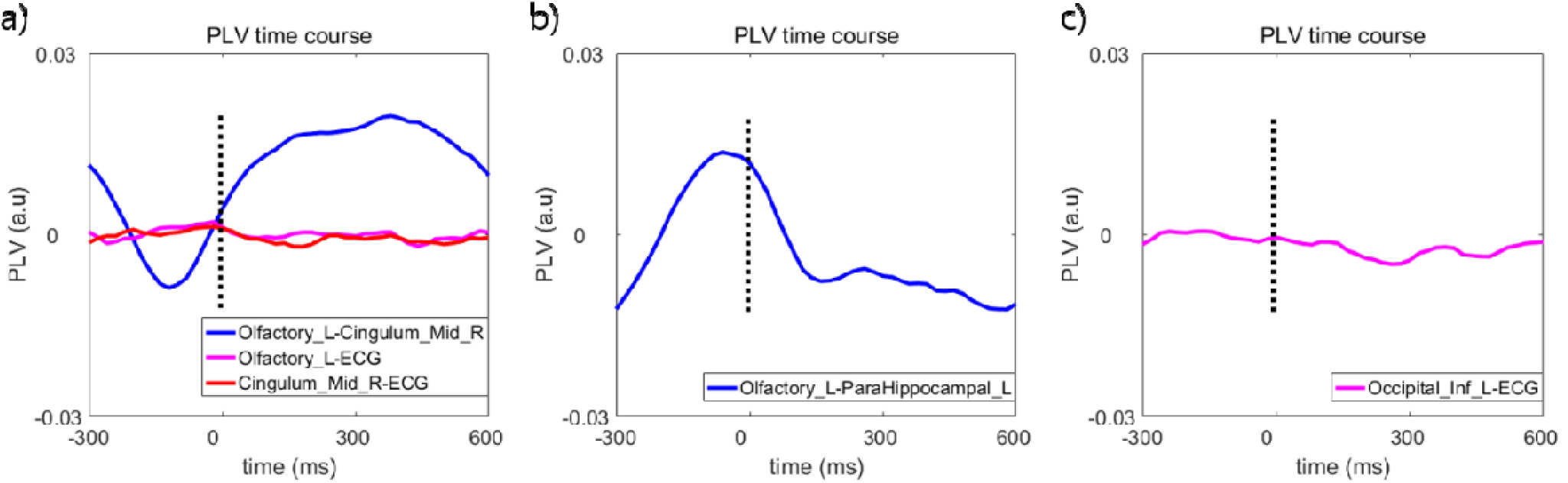
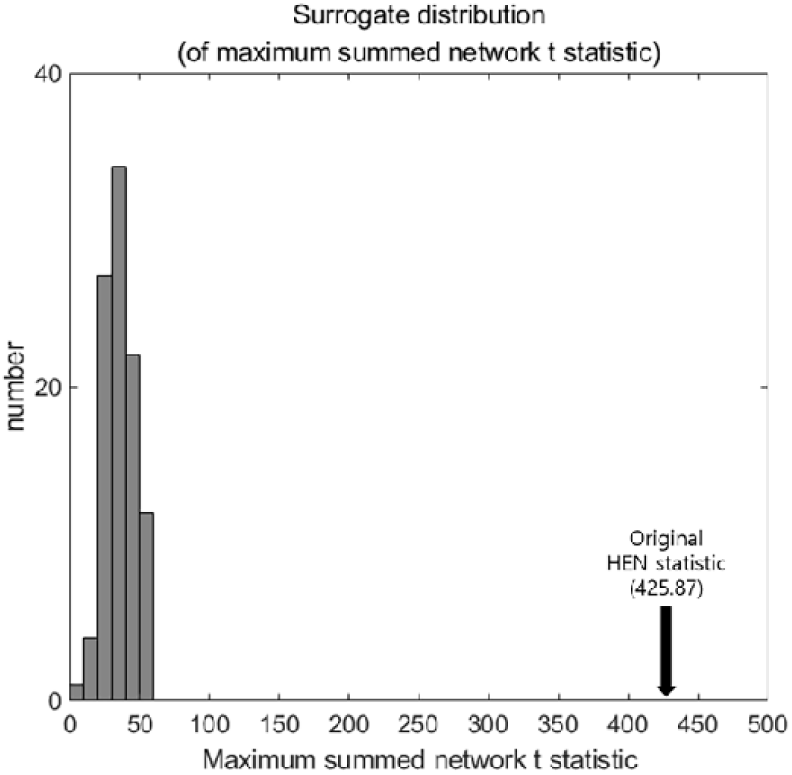
Representative plots of the changes in PLV over time. **a) Representative plot of the time courses of the theta PLVs between pairs of cortical regions and the theta PLVs between ECG signals and each cortical region.** The time courses of the theta PLV show that the phase synchronization between ‘Cingulum_Mid_R’ and ‘Olfactory_L’ significantly increases after a heartbeat, while the phase synchronization between ECG and ‘Cingulum_Mid_R’ and the phase synchronization between ECG-‘Olfactory_L’ does not change substantially. **b) Representative plot of the time course of the theta PLVs between cortical regions showing decreased PLVs after the heartbeat.** In contrast to the time courses of the theta PLVs between ‘Cingulum_Mid_R’ and ‘Olfactory_L’, phase synchronization between ‘Olfactory_L’ and ‘ParaHippocampal_L’ decreased over time. **c) Representative plot of the time course of the theta PLVs between ECG signals and cortical regions.** The phase synchronization between ECG signals and ‘Occipital_Inf_L’ did not show significant changes between the baseline and evoked periods. Note that in all PLV time courses plotted in the Figure 1, mean PLV of the baseline (300-100 ms before R-peak) was subtracted for a visualization purpose.

### No significant change in phase synchronization occurred between ECG signals and cortical regions

The paired t-tests (PLVs for ECG signals and cortical regions) comparing the responses between the evoked period (200-600 ms post R-peak) and baseline did not reveal a significant increase or decrease in PLVs between ECG signals and cortical regions in the theta band (the minimum p-value among 78 cortical regions was p = 0.246 (false discovery rate (FDR)- corrected) with t (86) = −2.86 in the ‘Occipital_Inf_L’, Figure 1c). The ‘Temporal_Pole_Mid_R’ region had the highest t-value, which was not significant (t (86) = 1.79, p = 0.076 (uncorrected)). If the electromagnetic field generated by cardiac contractile activity induced artificially increased phase synchronization between regions in the HEN compared with the baseline period, the ECG signal originating from the same electromagnetic field should have increased phase synchronization with cortical regions within the HEN. However, phase synchronization between cortical regions and ECG signals did not change in the evoked period compared to baseline, but phase synchronization between the regions in the HEN increased in the evoked period, indicating that the increased theta phase synchronization between cortical regions in the HEN was not caused by CFA (Figure 1a). Furthermore, the HEN was not likely caused by the pulse artefact (PA), which occurs when the sensors are influenced (moved) by vascular pulsation. First, in our study, we used MEG data, and MEG sensors do not directly contact the subject; thus, a vessel is unable to induce a pulsatile movement of sensors that causes PA. To our knowledge, no previous studies have reported a PA in MEG recordings. Second, according to a previous HER study using electrocorticography (ECoG) (Kern et al., 2013), if PA-induced artificial synchrony occurs between ECoG electrodes, the ECG and ECoG electrode likely display high phase synchronization (Kern et al., 2013), which was not observed in our results. By summarizing these results, the HEN we identified in theta frequency band was not caused by an artificial increase in phase synchronization induced by CFA or PA. While the theta-phase synchronization between ECG signals and cortical regions was not increased compared to the baseline, the CFA-induced increase in the phase synchronization compared with the baseline might exist in lower frequency bands, such as the delta band (0.5-4 Hz), because the cardiac contractile activity typically occurs at a rate of 60-100 beats per minute (BPM), which corresponds to a frequency of 1-1.67 Hz that belongs to the delta band. Similarly, in our data, subjects displayed a maximum heart rate of 88.84 BPM (∼ 1.48 Hz), thus the CFA or PA, particularly those induced by the T-wave, might have increased the artificial synchronization in the delta band.

### A surrogate R-peak analysis revealed that the effect of the HEN is time-locked to the heartbeat

No maximum network statistic in the surrogate R distribution was greater than our original network statistic of the HEN (network statistic = 425.87, Monte Carlo p < 0.01, Figure 2), indicating that the HEN identified in our study was most likely time-locked to the heartbeat.

**Figure 2.**
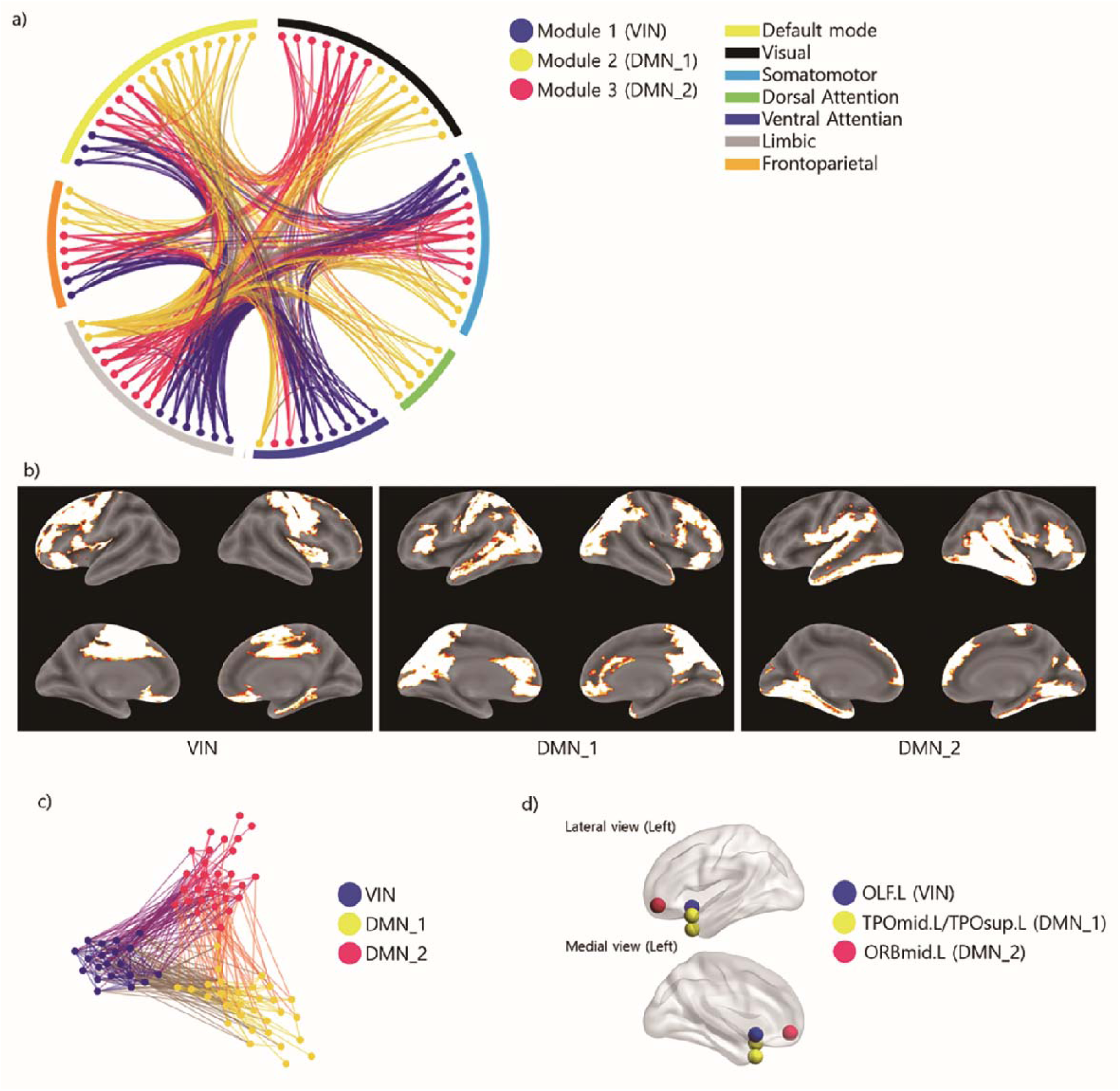
Surrogate R-peak distribution. The gray bars represent the distribution of the maximum summed network *t* statistic obtained from 100 surrogate R-peaks, and the black arrow indicates the original summed network *t* statistic.

### Modules of the static HEN

Based on the consensus partitioning results, the static HEN was partitioned into three modules (Figure 3b and Supplementary Table 2). Module 1 comprised the major visceromotor and interoceptive regions, including the bilateral middle cingulate cortex, insula, gyrus rectus, supplementary motor area, dorsolateral prefrontal cortices (dlPFC), right orbitofrontal cortices and right frontal operculum, which we named the visceromotor-interoceptive network (VIN). Module 2 comprised some DMN regions, including bilateral ventromedial prefrontal cortices (vmPFC), anterior/posterior cingulate cortices, the precuneus, and the left superior/middle temporal pole; this module was called default mode network 1 (DMN_1). Finally, Module 3 also contained regions of the DMN, including bilateral dorsomedial prefrontal cortices, lateral orbitofrontal cortices, lateral temporal cortices and the right superior/middle temporal pole; module 3 was called DMN_2. Notably, this module also contained areas of the ventral visual stream, including bilateral inferior occipital cortices, inferior temporal cortices and fusiform face areas (FFAs). Module 1 had the highest mean t-value for edges among the three modules, and the edges of Module 2 displayed higher mean t-values than those of Module 3 (Figure 3c, one-way ANOVA between the average edge t-values for the three modules, F (2, 225) = 22.21, p < 0.001, Module 1 > Module 2 in the post hoc two-sample t-test with t (158) = 3.87, p < 0.001 (Bonferroni-corrected), Module 2 > Module 3 in the post hoc two-sample t-test with t (138) = 2.73, p = 0.028 (Bonferroni-corrected)).

**Figure 3.**
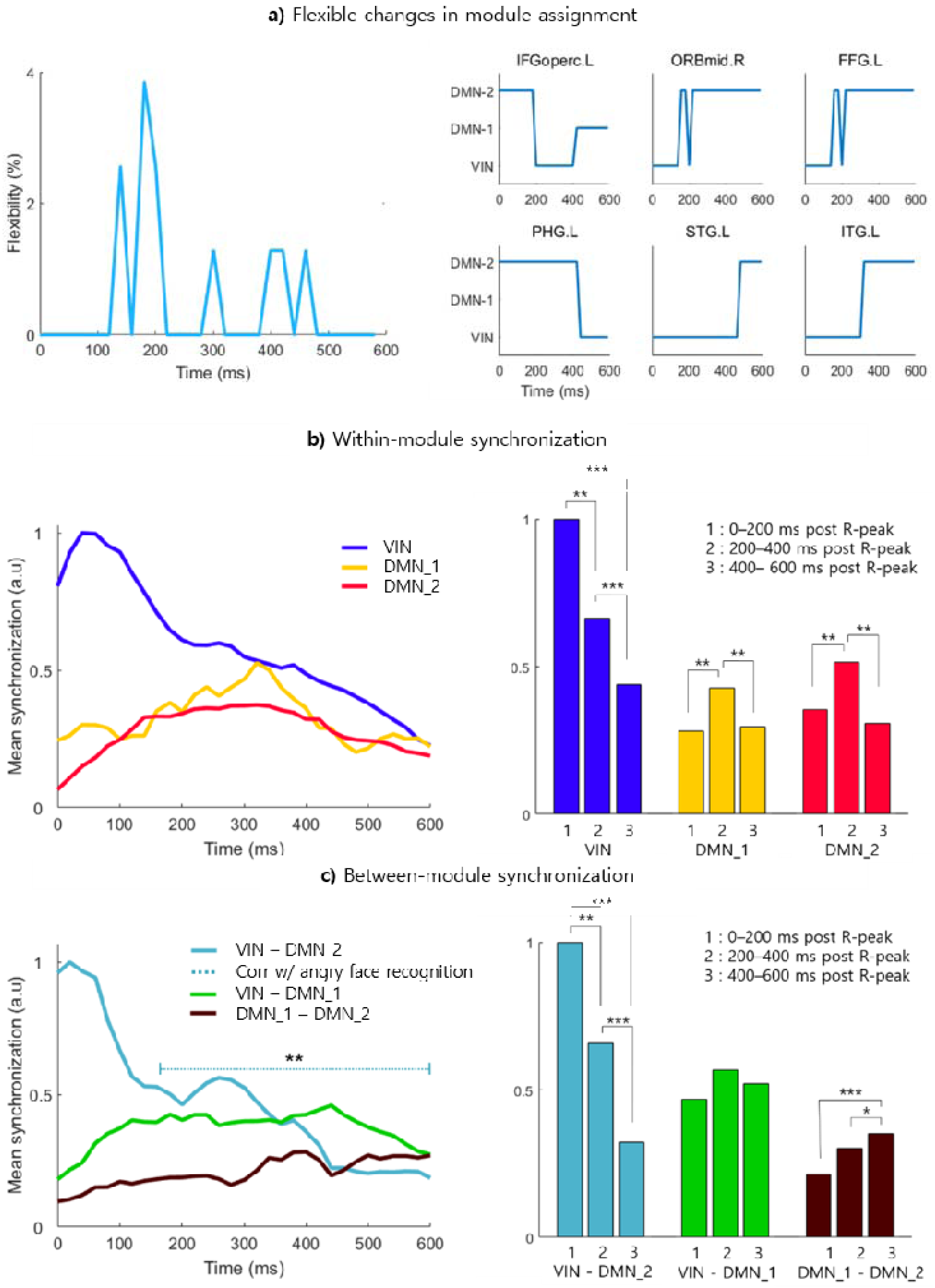
Structure of the HEN. **a) Relationship between the classical functional network and the HEN.** We calculated the spatial correlations between each region and the classical functional networks defined in previous studies (Thomas Yeo et al., 2011). Then, regions were assigned to the network with the maximum spatial correlation. Each classical functional network is represented as a colored arc, and the HEN module to which each region belongs is represented by the color of the node. Notably, most regions in the ventral attention network were VIN regions. This figure was generated using the connectogram (http://immersive.erc.monash.edu.au/neuromarvl/). **b) Surface map of three HEN modules.** The VIN included the major visceromotor and interoceptive regions, including the bilateral middle cingulate cortex and insula. DMN_1 was composed of part of the DMN regions bilateral ventromedial prefrontal cortices (vmPFC), anterior/posterior cingulate cortices, and precuneus. Finally, DMN_2 also contained regions of the DMN, including bilateral dorsomedial prefrontal cortices, lateral orbitofrontal cortices, lateral temporal cortices and the right superior/middle temporal pole. This figure was generated using bspmview software (dx.doi.org/10.5281/zenodo.168074). **c) Modular structure of the HEN.** Both the left and right figures show modular structures of the HEN. The lengths of the edges represent the strength of synchronization between regions according to the t-value. This figure shows larger synchronizations within regions of the VIN (blue), DMN_1 (orange), and DMN_2 (red) compared to the synchronization between modules. The synchronizations within the VIN regions (blue) were especially stronger than those within other modules. This figure was generated using the connectogram (http://immersive.erc.monash.edu.au/neuromarvl/). d) **Hubs of modules.** The hubs of each module were the left olfactory cortex (‘Olfactory_L’), left middle/superior temporal pole (‘TPOmid_L’ and ‘TPOsup_L’) and left lateral orbitofrontal cortex (‘ORBmid_L’), and these hubs are marked as white stars. This figure was created using BrainNet viewer (Xia, Wang, & He, 2013). A full list of the regions in each module is provided in the Supplementary Table 2.

### Hubs of the static HEN

Using graph theoretical measures, we identified the hubs of the HEN. Interestingly, the left middle temporal pole was determined to be a hub in every graph theoretical measure, including the z-P classification (Guimera & Amaral, 2005), nodal strength and betweenness centrality (Brandes, 2001) (Figure 3d and Table 1, this region had the highest within-module z-score, nodal strength and betweenness centrality among the 78 cortical regions). Moreover, each module had at least one connector hub that connects the nodes of different modules (Figure 3d and Table 1). The connector hubs of Modules 1, 2, and 3 were the left olfactory cortex, left middle/superior temporal pole and orbital part of the left middle frontal gyrus, respectively (Figure 3d and Table 1). Notably, all hubs are located in the left hemisphere.

**Table 1.** Node roles defined by graph theoretical measures (ordered by magnitude)

| **Connector Hub**<br>**(z, P)** | **High Strength Hub**<br>**(Nodal strength)** | **High BC Hub**<br>**(BC)** |
| --- | --- | --- |
| Temporal_Pole_Mid_L<br>(3.95, 0.58) | Temporal_Pole_Mid_L<br>(0.16) | Temporal_Pole_Mid_L<br>(2176) |
| Frontal_Mid_Orb_L<br>(3.47, 0.5) | Temporal_Pole_Sup_L<br>(0.14) | Frontal_Mid_Orb_L<br>(1116) |
| Olfactory_L<br>(2.98, 0.54) | Olfactory_L<br>(0.13) | Olfactory_L<br>(596) |
| Temporal_Pole_Sup_L<br>(2.65, 0.58) | Frontal_Inf_Orb_L<br>(0.11) | Cingulum_Mid_L<br>(512) |
|  | Temporal_Inf_L<br>(0.09) | Temporal_Inf_L<br>(508) |
|  | Frontal_Mid_Orb_L<br>(0.09) | Frontal_Inf_Orb_L<br>(478) |

**BC: Betweenness centrality**

### Modular structure of the dynamic HEN

Consensus partitioning of the multilayer network revealed three modules, and these results were similar to the consensus partitioning results from the static modularity analysis (Supplementary Table 3 and Supplementary Figure 1). Most regions of each module identified in the static modularity analysis were also present in modules identified in the dynamic modularity analysis. Notably, the connector hubs of each module in the static modularity analysis were present in the same module in the dynamic modularity analysis at all layers (Supplementary Table 3). However, because we included the time window before the 200 ms post R-peak, which was not used in the static or dynamic modularity analyses, the link between consecutive time points was also considered, and each module exhibited a slightly different composition and size. Notably, the VIN in the dynamic modularity analysis was larger than the same network in the static modularity analysis, potentially indicating that the VIN was larger in the period before the 200 ms post R-peak window than after the 200 ms post R-peak window.

### Changing modular composition of the dynamic HEN over time

The network structure of the HEN, which is defined as the composition of each module of the dynamic HEN, changed over time. For example, a the sizes of number of regions comprising the VIN slightly decreased from the 0 ms to 600 ms post R-peak values, while the sizes of DMN_1 and DMN_2 slightly increased over time (Supplementary Figure 1). An analysis of the flexibility of the entire brain over time qualitatively showed this trend (Figure 4a). The F_time was maximized at 180 ms post R-peak, which is the time closest to the 200 ms post R-peak, as we expected (3.85% of the total regions changed their community assignment at this time point, Figure 4a). Interestingly, all changes in community assignments at approximately 200 ms post R-peak occurred between the VIN and DMN_2 (Figure 4a).

**Figure 4.**
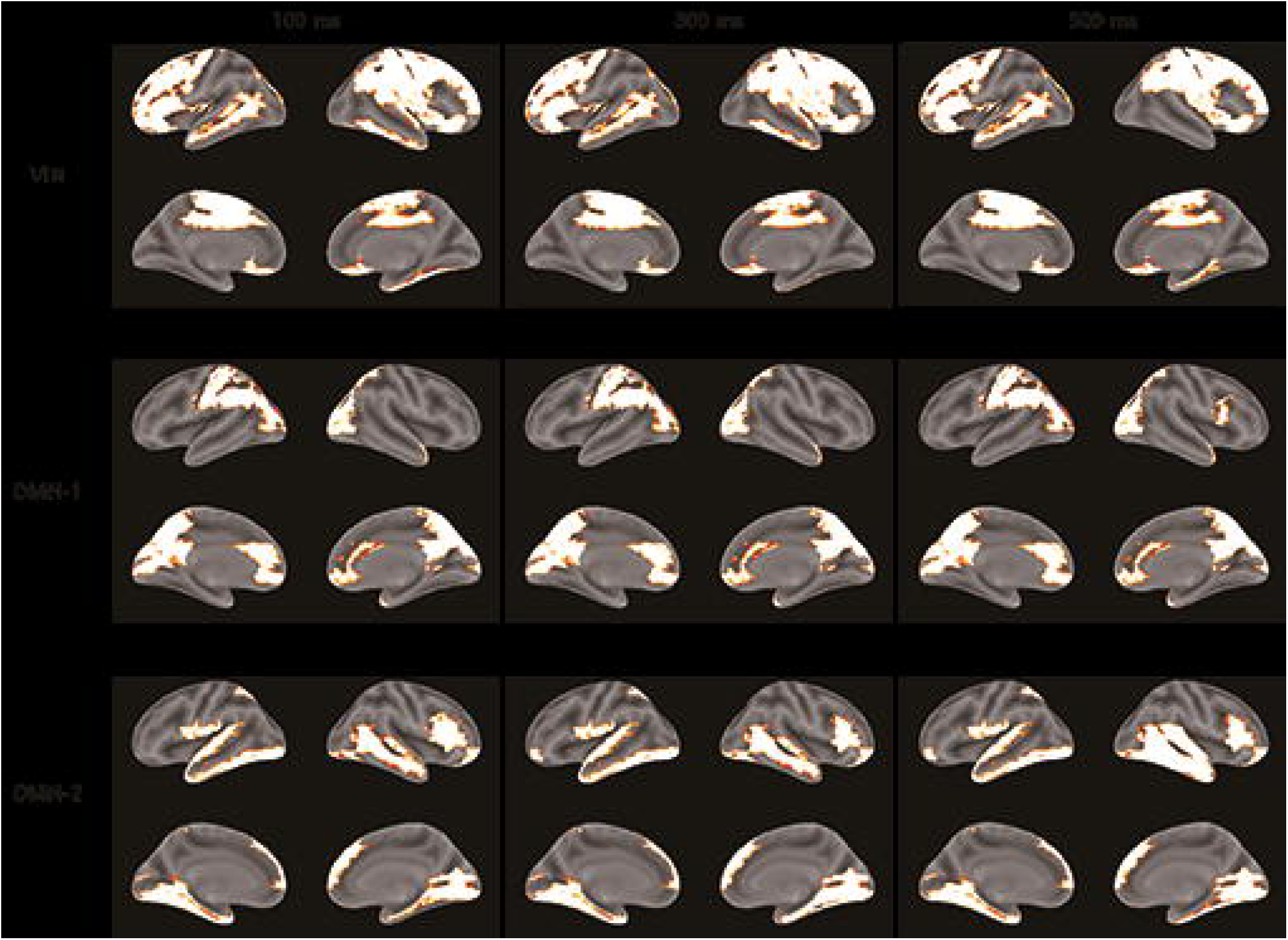
Dynamic changes in the HEN over time. **a). Flexible change in community assignment.** The left figure shows the whole-brain flexibility of the HEN over time. The flexibility of the HEN changed over time, and the maximum value was 3.85% at 180 ms post R-peak (left). The right figure shows the time courses of the community assignments in regions that changed module assignments at least one time. All changes in community assignments occurred between the VIN and DMN_2 except for ‘Frontal_Inf_Oper_L’, which changed the community assignment twice from DMN_2 to the VIN in the 180 ms post R- peak and from the VIN to DMN_1 in the 400 ms post R-peak (right). **b) Changes in within-module synchronization.** The within-module synchronization of each module changed over time. The within-module synchronization of the VIN (blue) was maximized initially and decreased over time, while the within-module synchronizations of both DMN_1 (pink) and DMN_2 (red) were maximum at the 320 ms post R-peak and decreased over time (left). This result was tested statistically by comparing the mean within-module synchronizations of the 0-200 ms, 200-400 ms, and 400-600 ms post R-peak time windows (right). Significant differences between time windows are marked. Synchronizations are scaled by the maximum synchronization for visualization. **c) Changes in between-module synchronization.** The between-module synchronization also changed over time. The synchronization between the VIN (blue) and DMN_2 (red) was maximized initially and decreased over time, while the synchronization between DMN_1 (pink) and DMN_2 (red) continually increased. (left). This result was tested statistically by comparing the mean between-module synchronizations of the 0-200 ms, 200-400 ms, and 400-600 ms post R-peak time windows (right). Significant differences between the time windows are marked. The dashed cyan line indicates the time window that showed a significant correlation between the angry face recognition score and synchronization between the VIN and DMN_2. * p < 0.05, ** p < 0.01, *** p < 0.001. ORBmid.R: right middle frontal gyrus, orbital part; INFoperc.L: left inferior operculum; PHG.L: left parahippocampal gyrus; FFG.L: left fusiform gyrus, STG.L: left superior temporal gyrus; ITG.L: left inferior temporal gyrus. Corr w/ angry face recognition: Correlation with the angry face recognition score.

### Changes among interactions within and between modules of the dynamic HEN over time

According to the results of an analysis of within-module synchronization, the synchronizations within all modules differed significantly between time windows (VIN: F (2, 172) = 18.33, p < 0.001); DMN_1: F (2, 172) = 7.422, p < 0.001; DMN_2: F (2, 172) =11.607, p < 0.001, Figure 4b). Post hoc paired t-tests revealed decreases in the within-module synchronizations of the VIN over time (0-200 ms > 200-400 ms, t (86) = 3.27 p = 0.005, 200-400 ms > 400-600 ms, t (86) = 3.77 p < 0.001, both p-values of paired t-tests were subjected to the Bonferroni correction). Moreover, the within-module synchronizations of DMN_1 and DMN_2 were significantly larger in the 200-400 ms time window than in the 0-200 ms and 400-600 ms time windows (DMN_1: 0-200 ms < 200-400 ms, t (86) = 3.21 p = 0.006, 200-400 ms > 400-600 ms, t (86) = 3.16 p = 0.006; DMN_2: 0-200 ms < 200-400 ms, t (86) =3.48 p < 0.001, 200-400 ms > 400-600 ms, t (86) = 4.42 p < 0.001, all p-values of paired t-tests were subjected to the Bonferroni correction). In the analysis of between-module synchronization, the synchronization between the VIN and DMN_2 was initially maximized and decreased over time (Figure 4c). Repeated measures ANOVAs of the between-module synchronizations of the VIN and DMN_2 between the 3 time windows revealed significant differences (F (2,172) = 26.38, p < 0.001, Figure 4c). Post hoc t-tests detected significantly greater synchronization between the VIN and DMN_2 in the 0-200 ms window than in the 200-400 ms (t (86) = 3.05, p = 0.009, Bonferroni-corrected) and 400-600 ms time windows (t(86) = 6.34, p < 0.001, Bonferroni-corrected). However, the synchronization between DMN_1 and DMN_2 increased after the 200 ms post R-peak compared to before the 200 ms post R-peak. Repeated measures ANOVAs between time windows showed a significant difference (F (2,172) = 9.36, p < 0.001, Figure 4c), and post hoc t-tests showed significantly greater synchronizations of DMN_1 and DMN_2 in the 200-400 ms post R-peak than in the 0-200 ms R-peak (t (86) = 2.84, p = 0.017, Bonferroni-corrected) and 400-600 ms post R-peak (t (86) = 3.76, p < 0.001, Bonferroni-corrected). However, significant increases in the 400-600 ms post R-peak were not observed compared to the 200-400 ms post R-peak (t (86)= 1.92, p = 0.058). Finally, significant differences in synchronization between DMN_1 and the VIN were not observed over time (F (2,172) = 1.66, p = 0.194).

### The between-module synchronization between the VIN and DMN_2 correlated with the ability to recognize angry faces

In the cluster-based permutation correlation analysis between the anger recognition score from the Penn Emotion Recognition Test and between-module synchronization, the synchronization between the VIN and DMN_2 in the 160-600 ms post R-peak negatively correlated with the angry face recognition score (Monte Carlo p = 0.005, Figure 4c - left panel); no other within/between-module synchronizations showed significant correlations. In addition, within/between-module synchronizations were not significantly correlated with the fearful face recognition score (minimum Monte Carlo p = 0.256).

## Discussion

In the resting state, our brain receives cardiac afferent signals, and previous studies showed a regional modulation of the HER under interoception-related task conditions. As shown in the present study, a heartbeat evokes signals from a network consisting of increased theta phase synchronization between cortical regions, which we called the HEN. The HEN was composed of the VIN and two DMN modules. Every module had at least one connector hub. According to the results of an analysis of the dynamic HEN, synchronizations within and between modules changed over time, and the composition of modules also changed. This change was most notable around the 200 ms post R-peak, as reflected by the maximum flexibility at that time point. Finally, the relationship between the HEN and interoception-related function was revealed based on a significant correlation between the between-module synchronization and the angry face recognition ability.

An important step in proving the existence of the HEN was showing whether it represents an actual increase in neural synchronization or artificially increased synchronization induced by CFA of cardiac contractile activity or the PA. We refuted the possibility of artificially increased synchronization by analyzing the increase in ECG-cortical region phase synchronization. Therefore, we have revealed the existence of the HEN, which is likely to be composed of increased neural phase synchronization. However, because the artefact-induced increase in the phase synchronization compared with the baseline might exist in a lower frequency band, such as the delta band, we suggest that in the case of the MEG, an investigation of the HEN in the frequency ranges covering the theta and higher frequency bands would be more reliable because these bands are unlikely to be influenced by CFA or PA, while an investigation of the HEN in the delta band and lower frequency bands would be less reliable because artefacts and the HEN would be difficult to discriminate in these frequency bands.

The existence of the HEN showed that the processing of heartbeat information occurs at the network level, rather than at the level of a single region. Furthermore, the HEN was segregated into three modules, suggesting that these modules might support different kind of heartbeat-related information processing. The first module, which we called the VIN, contained interoceptive regions that receive an ascending visceral signal and visceromotor regions, indicating that the VIN directly interacts with the heart. Moreover, the synchronizations induced by the heartbeat within the VIN were the strongest among the modules. Thus, the major network-level heartbeat processing is likely to occur within the VIN and could be used to understand previous task-related regional HER modulations at the network level. For example, most of the regions displaying an HER modulation by heartbeat attention (Canales-Johnson et al., 2015; Pollatos et al., 2005) and emotion processing (Couto et al., 2015) occurred in regions of the VIN. Based on these findings, HER modulation in these regions is likely a part of network-level processing within the VIN, which includes an interaction between regions, and this hypothesis could be clarified by analyzing the HEN during these tasks. Furthermore, a recent resting-state HER study reported phase resetting by the heartbeat within regions including the insula and operculum, which are included in the VIN, and the strongest effect was observed in the theta band (Park et al., 2017). We postulate that this phase resetting within multiple regions is potentially related to a network-level synchronization within the VIN.

DMN_1 and DMN_2 contained regions of the default mode network, which showed correlated activity in the numerous previous studies (Fox et al., 2005). However, researchers have not determined whether these regions show network-level synchronization during heartbeat processing. In previous HER studies using the thought sampling task, the HERs of DMN regions correlated with self-related thinking (Babo-Rebelo, Richter, et al., 2016; Babo-Rebelo, Wolpert, et al., 2016), and we suggest that these region-level HER modulations induced by self-related thinking are also related to a network-level interaction within DMN_1/DMN_2 that could be investigated by analyzing the HEN during this task.

We identified hubs of the HEN. Because visceral signals, including the heartbeat, initially enters the insula in the VIN (Craig, 2009), the heartbeat information processing occurring in DMN_1/DMN_2 is likely to be induced by an interaction with the VIN, rather than by a direct interaction with the heartbeat. We propose that the connector hubs play an important role in these interactions between the VIN and DMN_1/DMN_2. In particular, the temporal pole, which was the most important HEN hub, was also shown to function as a connector hub of the unified allostatic-interoceptive network composed of the salience network and default mode network, in a previous study (Kleckner et al., 2017). Considering that the salience network investigated in that study is very similar to the VIN, the result of that study also supports our hypothesis.

In addition to the static HEN, the analysis of the dynamic HEN showed changes in the structure of the HEN over time, and these changes were most prominent at approximately 200 ms post R-peak, which was represented by the maximal flexibility, suggesting that the most critical event that changes the HEN structure is the entry of the heartbeat into the cortex. Furthermore, while one might expect that the synchronization of the VIN would increase after 200 ms post R-peak to process the bottom-up heartbeat signal, our results revealed the greatest synchronization within the VIN in the 0-200 ms post R-peak time bin and decreased over time. This result is counterintuitive from the perspective of bottom-up sensory processing. However, we suggest that this result is consistent with the embodied interoceptive predictive coding (EPIC) model (Barrett & Simmons, 2015). The EPIC model suggests that agranular visceromotor cortices, such as the anterior insula and cingulate cortices, send signals resulting from allostatic visceromotor activity to the body and simultaneously issue interoceptive predictions to the granular interoceptive cortices regarding the homeostatic signal, and the interoceptive prediction is updated by an interoceptive prediction error (Barrett & Simmons, 2015). In a recent study, the salience network and the DMN were reported to support a unified allostatic-interoceptive system, which is essential in the EPIC model (Kleckner et al., 2017). The results of this study are similar to the HEN modules, as the VIN and the DMN_1/DMN_2 identified in our study correspond to the saliency network and DMN described in the previous study, respectively. From this perspective, the maximum synchronization of the VIN observed before the heartbeat enters the CNS might represent an interoceptive prediction of the heartbeat, and if an upcoming heartbeat does not make an interoceptive prediction error that corresponds to the resting state, the synchronization within the VIN would not increase, as evidenced by the within-module synchronization of the VIN. However, if an interoceptive prediction error occurs through an unexpected interoceptive event such as emotional arousal or by an increased the precision of the bottom-up heartbeat signal that increases the precision-weighted interoceptive prediction error (e.g., heartbeat attention task) (Petzschner et al., 2019), the synchronization would increase after the heartbeat, consistent with the results of previous HER studies on emotion and heartbeat attention tasks showing activation within regions of the VIN (Couto et al., 2015; Pollatos et al., 2005). While the EPIC model supports our results, we did not report the existence of the interoceptive prediction error; thus, further experiments showing an interoceptive prediction error would help clarify the EPIC model account of the HEN.

A limitation of our study is that while interoceptive processing typically includes subcortical regions, such as the amygdala (Kleckner et al., 2017), we used only cortical regions to construct the HEN. A HEN including subcortical regions might contain different structures and hubs than those reported here. Therefore, future studies on HENs including subcortical regions using deep source imaging MEG techniques are needed.

In conclusion, we first determined the existence of the theta oscillatory HEN that was partitioned into three modules consisting of the VIN and two DMNs, where the suggested role of the VIN was the network-level processing of the heartbeat and DMNs receive this processed information through an interaction with VIN via the connector hubs. Furthermore, the change in the structure of the HEN observed before and after the time the heartbeat enters the CNS, including the strongest synchronization of the VIN before this time point, suggests the a change in the HEN structure from prediction to the bottom-up processing of the heartbeat, consistent with the EPIC model.

Our results provide a framework that potentially contextualizes previous studies of the regional level HER modulation from a network perspective, which should be investigated in further studies analyzing the HEN during tasks. Furthermore, similar to the previous studies showing that a RSN is related to task performance (van Dam et al., 2015), the HEN was related to the emotion recognition score. Therefore, an interesting further study would be focused on investigating the relationships between other interoceptive functions and the HEN. Finally, investigations of the HEN in patients with a psychiatric disease related to interoception, such as an anxiety disorder (Paulus & Stein, 2010), might also improve our understanding of the pathophysiology.

## Figure legends

**Supplementary Table 1.** List of 78 cortical regions used in the analysis.

| **Region name (AAL)** | **Abbreviation** | **Region name (AAL)** | **Abbreviation** |
| --- | --- | --- | --- |
| 'Precentral_L' | PreCG.L | 'Calcarine_R' | CAL.R |
| 'Precentral_R' | PreCG.R | 'Cuneus_L' | CUN.L |
| 'Frontal_Sup_L' | SFGdor.L | 'Cuneus_R' | CUN.R |
| 'Frontal_Sup_R' | SFGdor.R | 'Lingual_L' | LING.L |
| 'Frontal_Sup_Orb_L' | ORBsup.L | 'Lingual_R' | LING.R |
| 'Frontal_Sup_Orb_R' | ORBsup.R | 'Occipital_Sup_L' | SOG.L |
| 'Frontal_Mid_L' | MFG.L | 'Occipital_Sup_R' | SOG.R |
| 'Frontal_Mid_R' | MFG.R | 'Occipital_Mid_L' | MOG.L |
| 'Frontal_Mid_Orb_L' | ORBmid.L | 'Occipital_Mid_R' | MOG.R |
| 'Frontal_Mid_Orb_R' | ORBmid.R | 'Occipital_Inf_L' | IOG.L |
| 'Frontal_Inf_Oper_L' | IFGoperc.L | 'Occipital_Inf_R' | IOG.R |
| 'Frontal_Inf_Oper_R' | IFGoperc.R | 'Fusiform_L' | FFG.L |
| 'Frontal_Inf_Tri_L' | IFGtriang.L | 'Fusiform_R' | FFG.R |
| 'Frontal_Inf_Tri_R' | IFGtriang.R | 'Postcentral_L' | PoCG.L |
| 'Frontal_Inf_Orb_L' | ORBinf.L | 'Postcentral_R' | PoCG.R |
| 'Frontal_Inf_Orb_R' | ORBinf.R | 'Parietal_Sup_L' | SPG.L |
| 'Rolandic_Oper_L' | ROL.L | 'Parietal_Sup_R' | SPG.R |
| 'Rolandic_Oper_R' | ROL.R | 'Parietal_Inf_L' | IPL.L |
| 'Supp_Motor_Area_L' | SMA.L | 'Parietal_Inf_R' | IPL.R |
| 'Supp_Motor_Area_R' | SMA.R | 'SupraMarginal_L' | SMG.L |
| 'Olfactory_L' | OLF.L | 'SupraMarginal_R' | SMG.R |
| 'Olfactory_R' | OLF.R | 'Angular_L' | ANG.L |
| 'Frontal_Sup_Medial_L' | SFGmed.L | 'Angular_R' | ANG.R |
| 'Frontal_Sup_Medial_R' | SFGmed.R | 'Precuneus_L' | PCUN.L |
| 'Frontal_Med_Orb_L' | ORBsupmed.L | 'Precuneus_R' | PCUN.R |
| 'Frontal_Med_Orb_R' | ORBsupmed.R | 'Paracentral_Lobule_L' | PCL.L |
| 'Rectus_L' | REC.L | 'Paracentral_Lobule_R' | PCL.R |
| 'Rectus_R' | REC.R | 'Heschl_L' | HES.L |
| 'Insula_L' | INS.L | 'Heschl_R' | HES.R |
| 'Insula_R' | INS.R | 'Temporal_Sup_L' | STG.L |
| 'Cingulum_Ant_L' | ACG.L | 'Temporal_Sup_R' | STG.R |
| 'Cingulum_Ant_R' | ACG.R | 'Temporal_Pole_Sup_L' | TPOsup.L |
| 'Cingulum_Mid_L' | DCG.L | 'Temporal_Pole_Sup_R' | TPOsup.R |
| 'Cingulum_Mid_R' | DCG.R | 'Temporal_Mid_L' | MTG.L |
| 'Cingulum_Post_L' | PCG.L | 'Temporal_Mid_R' | MTG.R |
| 'Cingulum_Post_R' | PCG.R | 'Temporal_Pole_Mid_L' | TPOmid.L |
| 'ParaHippocampal_L' | PHG.L | 'Temporal_Pole_Mid_R' | TPOmid.R |
| 'ParaHippocampal_R' | PHG.R | 'Temporal_Inf_L' | ITG.L |
| 'Calcarine_L' | CAL.L | 'Temporal_Inf_R' | ITG.R |

**R: right; L: left; Sup: superior; Mid: middle; Inf: inferior; Ant: anterior; Post: posterior; Med: medial;**

**Supplementary Table 2.** Regions in each module (static modularity analysis)

| VIN | DMN-1 | DMN-2 |
| --- | --- | --- |
| 'Precentral_L' | 'Frontal_Mid_L' | 'Frontal_Sup_Orb_L' |
| 'Precentral_R' | 'Frontal_Inf_Oper_L' | 'Frontal_Mid_Orb_L' |
| 'Frontal_Sup_L' | 'Frontal_Inf_Tri_R' | 'Frontal_Mid_Orb_R' |
| 'Frontal_Sup_R' | 'Frontal_Inf_Orb_L' | 'Frontal_Inf_Tri_L' |
| 'Frontal_Sup_Orb_R' | 'Frontal_Med_Orb_L' | 'Rolandic_Oper_L' |
| 'Frontal_Mid_R' | 'Frontal_Med_Orb_R' | 'Rolandic_Oper_R' |
| 'Frontal_Inf_Oper_R' | 'Cingulum_Ant_L' | 'Frontal_Sup_Medial_L' |
| 'Frontal_Inf_Orb_R' | 'Cingulum_Ant_R' | 'Frontal_Sup_Medial_R' |
| 'Supp_Motor_Area_L' | 'Cingulum_Post_L' | 'ParaHippocampal_R' |
| 'Supp_Motor_Area_R' | 'Cingulum_Post_R' | 'Calcarine_L' |
| 'Olfactory_L' | 'Calcarine_R' | 'Lingual_L' |
| 'Olfactory_R' | 'Cuneus_L' | 'Lingual_R' |
| 'Rectus_L' | 'Cuneus_R' | 'Occipital_Inf_L' |
| 'Rectus_R' | 'Occipital_Sup_L' | 'Occipital_Inf_R' |
| 'Insula_L' | 'Occipital_Sup_R' | 'Fusiform_L' |
| 'Insula_R' | 'Occipital_Mid_L' | 'Fusiform_R' |
| 'Cingulum_Mid_L' | 'Occipital_Mid_R' | 'SupraMarginal_L' |
| 'Cingulum_Mid_R' | 'Postcentral_R' | 'SupraMarginal_R' |
| 'ParaHippocampal_L' | 'Parietal_Sup_L' | 'Angular_R' |
| 'Postcentral_L' | 'Parietal_Sup_R' | 'Paracentral_Lobule_L' |
|  | 'Parietal_Inf_L' | 'Heschl_R' |
|  | 'Parietal_Inf_R' | 'Temporal_Sup_L' |
|  | 'Angular_L' | 'Temporal_Sup_R' |
|  | 'Precuneus_L' | 'Temporal_Pole_Sup_R' |
|  | 'Precuneus_R' | 'Temporal_Mid_L' |
|  | 'Paracentral_Lobule_R' | 'Temporal_Pole_Mid_R' |
|  | 'Heschl_L' | 'Temporal_Inf_L' |
|  | 'Temporal_Pole_Sup_L' | 'Temporal_Inf_R' |
|  | 'Temporal_Mid_R' |  |
|  | 'Temporal_Pole_Mid_L' |  |

**R: right; L: left; Sup: superior; Mid: middle; Inf: inferior; Ant: anterior; Post: posterior; Med: medial;**

**Supplementary Table 3.**
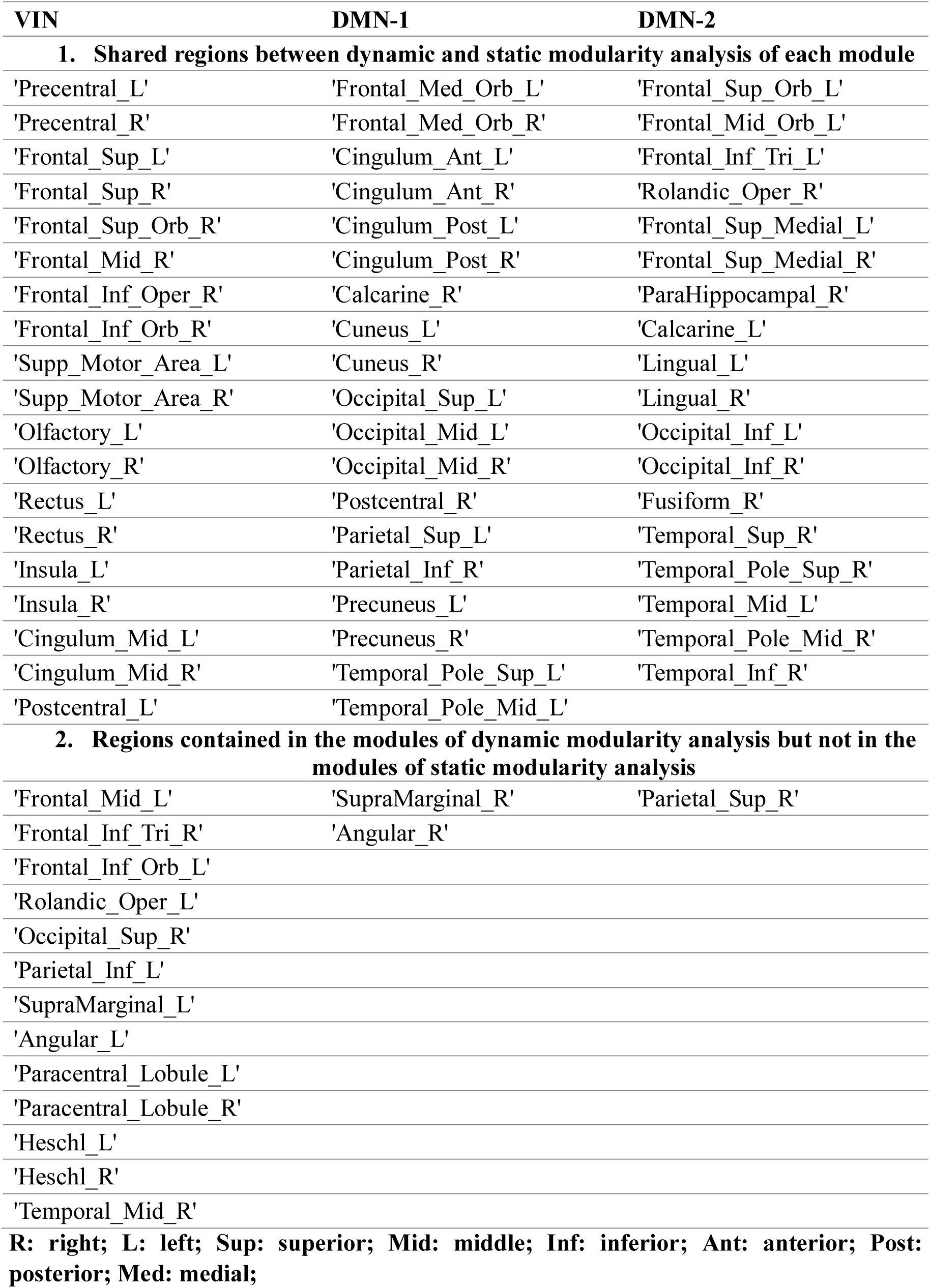
Regions that are consistently contained in the modules for every time points of dynamic modularity analysis.

## Acknowledgments

Data were provided by the Human Connectome Project of the WU-Minn Consortium (principal investigators: David Van Essen and Kamil Ugurbil; 1U54MH091657), which was funded by the 16 NIH Institutes and Centers that support the NIH Blueprint for Neuroscience Research and the McDonnell Center for Systems Neuroscience at Washington University. This research was supported by the Brain Research Program through the National Research Foundation of Korea (NRF) funded by the Ministry of Science & ICT (NRF - 2016M3C7A1914448 and NRF - 2017M3C7A1031331 to B.J.). This manuscript was edited for English language by American Journal Experts (AJE). The authors wish to acknowledge Hyeong-Dong Park for helpful comments and discussions on manuscript.

